# Mathematical deconvolution of CAR T-cell proliferation and exhaustion from real-time killing assay data

**DOI:** 10.1101/786020

**Authors:** Prativa Sahoo, Xin Yang, Daniel Abler, Davide Maestrini, Vikram Adhikarla, David Frankhouser, Heyrim Cho, Vanessa Machuca, Dongrui Wang, Michael Barish, Margarita Gutova, Sergio Branciamore, Christine E. Brown, Russell C. Rockne

## Abstract

Chimeric antigen receptor (CAR) T-cell therapy has shown promise in the treatment of hematological cancers and is currently being investigated for solid tumors including high-grade glioma brain tumors. There is a desperate need to quantitatively study the factors that contribute to the efficacy of CAR T-cell therapy in solid tumors. In this work we use a mathematical model of predator-prey dynamics to explore the kinetics of CAR T-cell killing in glioma: the Chimeric Antigen Receptor t-cell treatment Response in GliOma (CARRGO) model. The model includes rates of cancer cell proliferation, CAR T-cell killing, CAR T-cell proliferation and exhaustion, and CAR T-cell persistence. We use patient-derived and engineered cancer cell lines with an in vitro real-time cell analyzer to parameterize the CARRGO model. We observe that CAR T-cell dose correlates inversely with the killing rate and correlates directly with the net rate of proliferation and exhaustion. This suggests that at a lower dose of CAR T-cells, individual T-cells kill more cancer cells but become more exhausted as compared to higher doses. Furthermore, the exhaustion rate was observed to increase significantly with tumor growth rate and was dependent on level of antigen expression. The CARRGO model highlights nonlinear dynamics involved in CAR T-cell therapy and provides novel insights into the kinetics of CAR T-cell killing. The model suggests that CAR T-cell treatment may be tailored to individual tumor characteristics including tumor growth rate and antigen level to maximize therapeutic benefit.

**Statement of Significance:** We utilize a mathematical model to deconvolute the nonlinear contributions of CAR T-cell proliferation and exhaustion to predict therapeutic efficacy and dependence on CAR T-cell dose and target antigen levels.

## Introduction

Chimeric antigen receptor (CAR) T-cell therapy is a targeted immunotherapy, demonstrating remarkable antitumor efficacy, particularly in the treatment of hematologic cancers [1,2]. CAR T-cell therapy is a specific type of immunotherapy where T-cells are genetically modified to recognize a tumor antigen thereby specifically redirecting T cell cytolytic activity. Inspired by the success of CAR T-cell therapy in liquid tumors, there has been a great interest in expanding the use of CAR T-cells for the treatment of solid tumors, such as glioblastoma (GBM), a highly aggressive form of primary brain cancer. Several clinical trials using CAR T-cells to treat GBM have been initiated all over the world [3–6]. At this early stage of clinical development, CAR T-cells offer much promise in solid tumors. However, the diversity of current clinical trials employing varying types of CARs for different solid tumors, target patient populations, and preconditioning regimes, presents a significant challenge in identifying which aspects of a given CAR T-cell treatment protocol are most critical for its effectiveness. An additional critical challenge for CAR T-cell therapy is the potential for transient-progression, where the cancer appears to progress before eventually responding to the treatment [7,8].

In order to address these challenges in CAR T-cell therapy for solid tumors, we endeavored to study the kinetics of CAR T-cell killing with an *in vitro* system and a mathematical model. Mathematical models are useful to describe, quantify, and predict multifaceted behavior of complex systems, such as interactions between cells. A mathematical model is a formalized method to hypothesize systems dynamics, and yield solutions that represent the system’s behavior under certain initial conditions. Mathematical models can be versatile and tested with clinical data which may be obtained *in vivo* from non-invasive imaging [9–11]. When additional information about the system becomes available, the model can be refined and adjusted accordingly. Many mathematical models have been developed to understand tumor progression to guide refinement of cancer therapy regimens [12–14]. As CAR T-cell therapy is a newly advanced treatment modality, relatively few studies have utilized computational modelling to understand and improve this cell-based therapy. Recently computational models have been developed to investigate cytokine release syndrome for toxicity management [15–17, effect of cytokine release syndrome on CAR T-cell proliferation [18], and mechanisms of CAR T-cell activation [19,20], dosing strategies[21]. However, it remains an open challenge how to use mathematical modeling to study and ultimately predict dynamics of CAR T-cell mediated cancer cell killing with respect to CAR T-cell dose, cancer cell proliferation, target antigen expression, and how these factors contribute to the overall effectiveness of CAR T-cell therapy.

Based upon our pre-clinical and clinical experience with our well-characterized IL13Rα2-targeted CAR T-cell therapy for recurrent glioblastoma [22,23], we have identified several factors which contribute to the effectiveness of CAR T-cells: rates of proliferation, exhaustion, persistence, and target cell killing. To study these various facets of CAR T-cell killing kinetics, we modeled the dynamics between cancer cells and CAR T-cells as a predator prey system with the CARRGO mathematical model: Chimeric Antigen Receptor t-cell treatment Response in GliOma. We used a real-time cell analyzer experimental system to estimate parameters of the mathematical model and then apply the model to *in vivo* human data with the long-term aim of developing a model which could be used to predict and eventually to optimize response.

## Methods

The **CAR** T-cell treatment **R**esponse to **G**li**O**ma (**CARRGO**) mathematical model is a variation on the classic Lotka-Volterra [24,25] predator-prey equations:

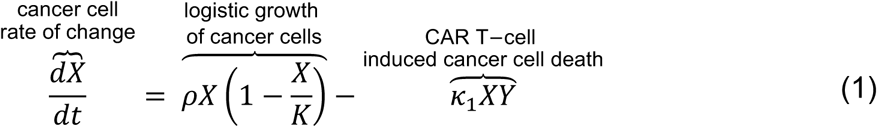

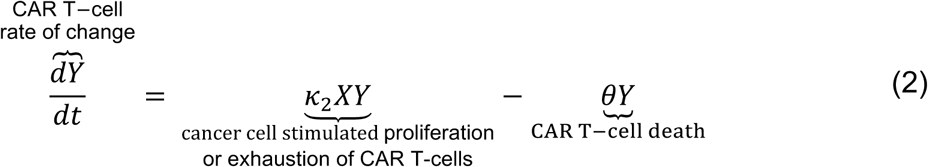

where *X* represents the density of cancer cells, *Y* is the density of CAR T-cells, *ρ* is the net growth rate of cancer cells, *K* is the cancer cell carrying capacity, *κ*_1_ is the killing rate of the CAR T-cells, *κ*_2_ is the net rate of proliferation including exhaustion or death of CAR T-cells when encountered by a cancer cell and *θ* is the natural death rate of CAR T-cells. The parameters *ρ, K, κ*_1_, *θ* are constants and assumed to be non-negative except for *κ*_2_ which can be either positive or negative. A positive value of *κ*_2_ indicates an increased rate of CAR T-cell proliferation when stimulated by interaction with a cancer cell. A negative value of *κ*_2_ indicates exhaustion or limited activation of CAR T-cells resulting from interaction with a cancer cell (**Table 1**). Exhaustion and hypoactivation of CAR T-cells are combined into a single value and are not modeled individually.

**Table 1:** CARRGO model parameters. All parameters are assumed to be non-negative except *κ* _2_ which may be positive or negative.

| Parameter | Description | Unit |
| --- | --- | --- |
| $\rho$ | Cancer cell net growth rate | day<sup>-1</sup> |
| $K$ | Carrying capacity | cell |
| $\kappa_1$ | CAR T-cell killing rate | day<sup>-1</sup> cell<sup>-1</sup> |
| $\kappa_2$ | Net rate of proliferation and exhaustion of T-cells when stimulated by cancer cells | day<sup>-1</sup> cell<sup>-1</sup> |
| $\theta$ | CAR T-cell net death (persistence) | day<sup>-1</sup> |

We chose to model the net number of cancer cells and simple interactions between cancer cells and CAR T-cells because the output data from the culture system is limited to cell number over time. We therefore are only able to infer dynamics at this scale and dimension (cells, time). Moreover, we performed a system identifiability analysis to demonstrate the parameters of model can be uniquely determined from the data in this experiment (see supplemental methods) [26–29]. Future studies may examine more complex dynamics and features such as modeling individual cell antigen levels, heterogeneity, resistant and sensitive sub populations, repeated treatments, etc. with other experimental designs which directly measure these features.

### Model assumptions

The CARRGO model treats cancer cell-CAR T-cell dynamics in this experimental condition as a closed predator-prey system. The model assumes 1) the populations are well mixed, 2) cancer cell growth is limited by space and nutrients (culture media) in the *in vitro* culture system and therefore grow logistically, 3) CAR T-cells kill cancer cells when they interact via the law of mass action, 4) the CAR T-cell killing rate does not explicitly assume a dependence on antigen density, 5) CAR T-cells may be stimulated to proliferate or to undergo loss of effector function—defined as exhaustion—upon contact with a cancer cell [30], and 6) the CAR T-cell death rate is independent of cancer cell density. We chose the Logistic growth model for the cancer cell population because the fixed growth rate and carrying capacity parameters were the biological quantities of interest when comparing CAR T-cell killing kinetics across cell lines. Witzel et al compared sigmoidal growth laws including Logistic, Gompertz, and Richards showed that all these models can be fit equally well to this form of experimental data [31]. Data supporting our model assumptions are given in supplemental material1 (**Fig. S1,S2**)

### Dynamical system analysis of the CARRGO model

Closed form solutions cannot be obtained for the relatively simple CARRGO model. To study the possible dynamics with the CARRGO model, we perform classical dynamical system analysis. Detailed mathematical analysis of this model can be found in several textbooks in dynamical systems [25,32]. In the interest of informing the reader, we briefly summarize the main points here. We begin by 1) scaling (non-dimensionalizing) the variables in the system and then 2) identify stationary points and classify their stability and finally 3) interpret the stationary points and system dynamics in terms of the initial numbers of cancer cells and CAR T-cells.

First, we scale the variables in the CARRGO model to obtain a model without physical units in order to study the intrinsic dynamics of the system. We scale time, the cancer cell and CAR T-cell populations as

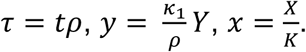

These variables are substituted into the CARRGO model (Eq.1,2) to obtain the scaled dimensionless system

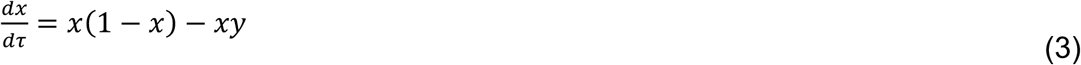

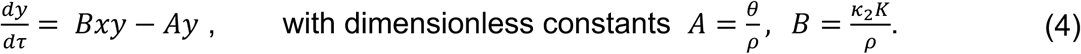

The steady-state solutions of this system are obtained by setting the time derivatives equal to zero (Eq. 3,4). The values of the dimensionless parameters A and B determine the dynamics of the system which may be represented as trajectories in a 2-dimensional phase-space of cancer cells and CAR T-cells (*x, y*). Three stationary points corresponding to steady-state solutions are denoted by *P*_*i*_ = (*x, y*) are: *P*_1_ = (0,0) where both the cancer cells and CAR T-cells are eliminated (zero), *P*_2_ = (1,0) where cancer cells population reaches the carrying capacity and CAR T-cells are eliminated, and a coexistence of both populations, 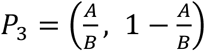. Three possible dynamics can result from this model depending on the values of A and B (**Fig.1**).

**Figure 1:**
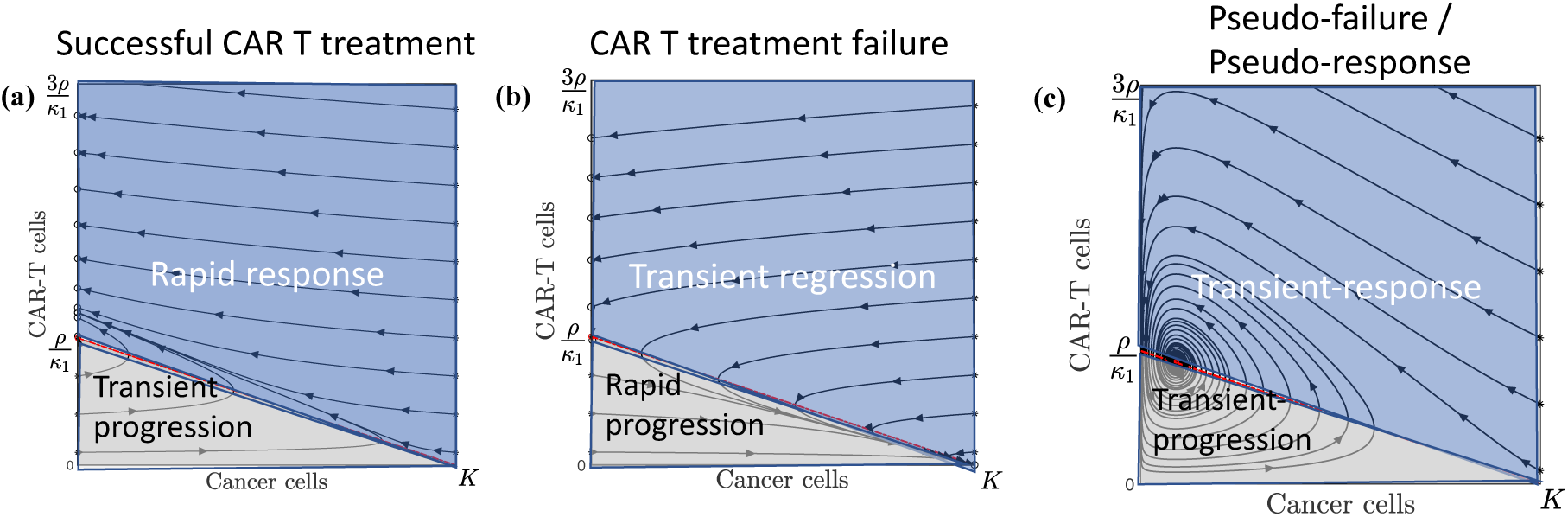
Possible dynamics from the CARRGO model. Dynamics are represented as trajectories in a 2-dimensional phase-space (*x, y*) = (cancer cells, CAR T-cells). (a) Case 1: Successful CAR T-cell treatment (A=0, B=0.2). This situation predicts all long-term dynamics to result in eradication of cancer cells with varying levels of residual CAR T-cells. (b) Case 2: CAR T treatment failure (A=0, B=−0.2). This situation predicts all long-term dynamics to result in cancer cells growing to carrying capacity and eventual elimination of CAR T-cells. (c) Case 3: Pseudo-failure/pseudo-response (A= 0.14, B=1.6). This situation predicts long-term coexistence of cancer cells and CAR T-cells, denoted P_3_ (red circle). In this situation, cancer cells and CAR T-cell populations increase, then decrease, then increase in an oscillatory manner. The dark blue regions shows cancer cell response and the light grey regions show cancer cell progression. We note that all 3 dynamics predicted by the CARRGO model include periods of transient increase or decrease in the cancer cell population, pointing to pseudo-progression of cancer, which a critical challenge in CAR T-cell treatment.

#### Case 1 Successful CAR T-cell treatment (*A* = 0, *B* > 0)

This situation occurs when the death rate of CAR T-cells is negligible (*θ* ≈ 0) relative to the proliferation rate of cancer cells (*ρ*), and the CAR T-cells are stimulated to proliferate when encountering a cancer cell (*κ*_2_ > 0). In this case the equilibrium points are when the cancer cells are eliminated with some remaining CAR T-cells (0, *y*) and when the cancer cells reach carrying capacity with no remaining CAR T-cells (K,0). Cancer cell elimination (0, *y*) is stable only if the CAR T-cell population is larger than the ratio (*y* > 1 ⇒ *Y* > *ρ*/*κ*_1_), that is to say if the product of the rate of CAR T-cell killing and the number of CAR T-cells is greater than the proliferation rate of the cancer cells. If the initial CAR T-cell population is below the line 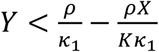, the CARRGO model predicts a *transient progression* of cancer cells before eventual response (grey region). The point (K,0) is an unstable repulsor state (**Fig.1a**).

#### Case 2 CAR T-cell treatment failure (*A* = 0, *B* < 0)

This situation occurs when the proliferation rate of CAR T-cell is less than the exhaustion rate of CAR T-cells due to interaction with cancer cells. In this case the fixed points are (0, *y*) and (K,0) are the same as in case 1. However, the point (K,0) is now a stable attractor state, corresponding to the cancer cells eventually growing to carrying-capacity and extinction of CAR T-cells. Again cancer cell elimination (0, *y*) is stable only if the CAR T-cell population is larger than the ratio (*Y* > *ρ*/*κ*_1_). If the initial CAR T-cell population is above the line 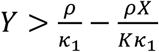, the CARRGO model predicts a *transient regression* of cancer cells before eventual rapid progression (grey region). This case is a failure of CAR T-cell treatment (**Fig.1b**).

#### Case 3 Pseudo-failure or pseudo-response (*A* > 0, *B* > 0)

In this situation, the third stationary point *P*_3_ corresponding to cancer cell and CAR T-cell coexistence lies in the first quadrant (positive numbers of cancer cells and CAR T-cells) only if *A* ≤ *B*. The point 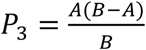 is then a stable sink (**Fig.1c**). This case results in a transient increase in cancer cells corresponding to tumor progression followed by a decrease in tumor cells corresponding to treatment response in an oscillating manner. The transient and oscillatory nature of these dynamics may be interpreted as a “pseudo”-failure and “pseudo”-response to the therapy. We note that cancer progression and treatment occur on finite and sometimes small timescales and therefore oscillatory dynamics may not be observed *in vivo* due to insufficient time to observe these changes.

### Cell lines

Low-passage primary brain tumor (PBT) lines were derived from GBM patients undergoing tumor resections at City of Hope as previously described [33,34]. Fibrosarcoma line HT1080 was obtained from the American Tissue Culture Collection (ATCC) and maintained according to recommendations. PBT030 endogenously expresses high level of IL13Rα2. HT1080 and PBT138 do not express IL13Rα2 and were lentivirally engineered to express varied levels based on different promoter strengths to investigate the relationship between killing kinetics and antigen expression level: High (>70%^+^) driven by the EF1α promoter, Medium (between 40%^+^-70%^+^) driven by the PGK promoter, Low (<20%^+^) driven by the attenuated PGK100 promoter [35,36]. These cell lines are denoted with H, M, L respectively e.x. HT1080-H. These tumor cell lines were selected because they differ in aggressiveness (proliferation rates) and antigen expression levels (endogenous or engineered).

Chimeric Antigen Receptor (CAR) T-cells were derived from healthy donor enriched CD62L+CD45RO+ central memory T cell population and lentivirally transduced with second-generation IL13Rα2-targeting CARs: IL13BBζ or IL1328ζ [33,34,37,38]. Transduced product was enriched for CAR and expanded in X-Vivo media with 10% FBS until 17 days in culture and cryopreserved. Non-transduced T cells expanded under the same condition was used as mock control.

### Experimental design

Real-time monitoring of cancer cell growth was performed by using xCELLigence cell analyzer system [39]. This system utilizes electrical impedance to non-invasively quantify adherent cell density with a dimensionless number referred to as cell-index (CI). The cell-index read-out from the machine is strongly positively correlated with the number of cells in the well (r^2^ > 0.9) and can be used as a linear measure of cell number [31]. We therefore report CARRGO parameter values in units CI which can be translated into units per cell based on the linear relation. Real-time cytotoxicity assay was performed using xCELLigence system in disposable 96 well E-Plates. Prior to seeding, tumor cells were enzymatically single-celled and seeded at 25 × 10^3^, 12.5 × 10^3^, or 2 × 10^3^ cells per well depending on the cell line. Cells were either left untreated (triplicates per cell line) or treated with CAR T-cells at effector to target ratios (E:T) of 1:5, 1:10, and 1:20. CAR T-cells were added to the wells about 24 hours after cancer cell seeding. Growth curves were recorded over 4 days with temporal resolution of 15 minutes (**Fig. 2**). Each cell line was treated with three IL13Rα2-targeted CAR T-cells: BBζ, 28ζ, and mock. At the end of the experiment, flow cytometry was performed to measure the residual CAR T-cells, cancer cells and IL13R*α*2 expression level. The details of cancer cell seeding, effector to target ratios used for the experiments are given in supplement material (**Table. S1**). The cancer cell dynamics of all the wells of 96 well E-plate for all cell lines are given in supplemental material (**Fig. S3**).

**Figure 2:**
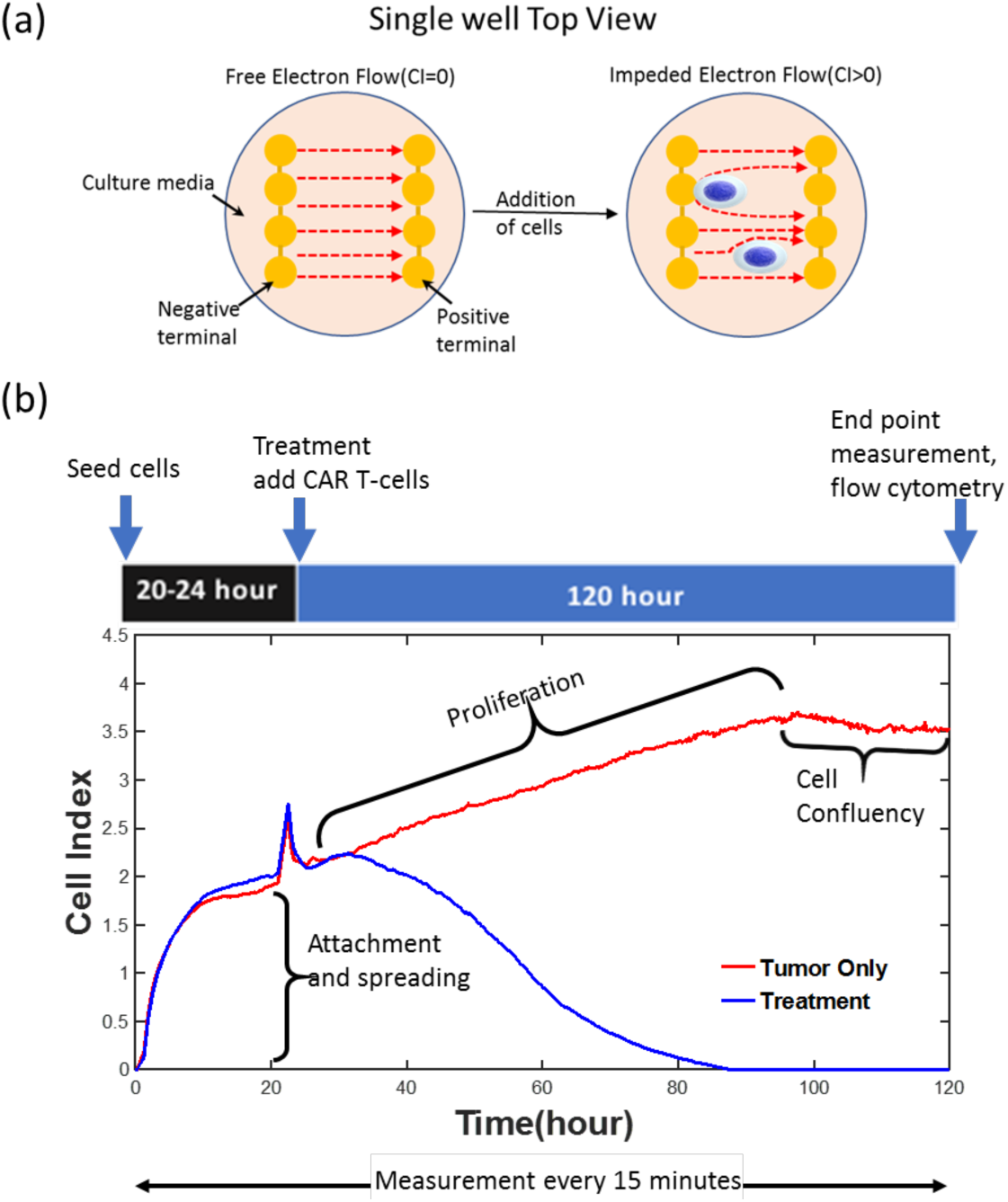
Schematic of experiment design and output data from the xCELLigence system. a) The xCELLigence system for real-time monitoring of cancer cell growth and response to CAR T-cell therapy. b) The output of xCELLigence is “Cell Index” (CI) which is calculated from changes in electrical impedance in the culture plate over time. Cell index is strongly correlated with the number of cells in the well/plate. This system results in distinct regimes of cell growth dynamics, including an attachment phase followed by proliferation and confluency. The proliferation growth regime was used to fit the CARRGO model and quantify dynamics of CAR T-cell killing of cancer cells (blue curve) as compared to untreated growth (red curve).

### CARRGO model fitting to experimental data

The first 24 hours of the time-series describes the process of cell attachment to the bottom of the plate (**Fig. 2**). The spatial process of cell adhesion and spreading in the well can be modeled as a reaction-diffusion process, described in supplement material (**Fig. S4**). Since we are interested in cell growth kinetics, we omitted the data from first 24 hours during the attachment process. The final time point is determined when the cells reach confluency which varies by cell line. Observed time of confluency for the PBT cell line was around 120 hours from the time of seeding while HT1080 reached confluency within 80 hours. The data points from CAR T-cell administration (24 hours after seeding) up to 80% of maximum CI value (confluency) were used for model fitting to estimate the parameters. At greater than 80% of the maximum CI, the linear relationship between CI and cell number no longer holds [31].

Cancer cell net growth rate *ρ* and carrying capacity *K* (Eq.1) were computed by fitting logistic growth to untreated cancer cell time series data. The CAR T-cell killing rate *κ*_1_, growth rate *κ*_2_, and death rate *θ* were computed by fitting the solution of the CARRGO model (Eqs.1,2) to treated cancer cell time series data by minimizing the root mean square error. All optimization computations were performed in MATLAB with *fmincon*.

### CARRGO model fitting to human data

A patient with recurrent glioma received CAR T-cells engineered for IL13R*α*2 and showed complete tumor regression, which was published as brief report by Brown et al in the New England Journal of Medicine in 2016 [33]. We retrospectively collected the magnetic resonance imaging (MRI) data of this patient and calculated tumor volumes to be used to fit the CARRGO model. Three lesions were selected using the lesion labeling reported in Brown et al [33]: lesions T6, T7 which responded to IL13Rα2 targeted CAR T-cells and lesion T9 which was a lesion that appeared later on which did not respond to the therapy. Tumor volumes for each lesion were estimated by manual segmentation of contrast-enhancing lesions from T1-weighted post-contrast MRIs. The number of cancer cells (CC) was estimated by calculating CC=[tumor volume (*μ*m^3^)]/[GBM cell size (*μ*m^3^)] with the average cell diameter assumed to be 20 *μ*m [40]. The volume of a spherical cell is then given by *V* = (4*π*/3)(10*μm*)^3^. These relationships were used to estimate the total number of cancer cells in a tumor volume. The tumor growth rate (*ρ* (1/time)) was computed from two subsequent imaging time points following the first appearance of the lesion on MRI. CAR T-cells were administered first directly into the tumor tissue and subsequently into the cerebrospinal fluid via the intraventricular injection. Because the CAR T-cells migrated to several tumor foci in the patient, we assumed a small fraction (5-10%) of the infused dose reached each individual lesion at each infusion. The CARRGO model was fit to the time-series MRI-derived tumor volume data by minimizing the root mean square error for MRIs before and during CAR T-cell treatment to compute the rates of CAR T-cell killing *κ*_1_, exhaustion *κ*_2_ and death θ.

## Results

### Model/data fitting to *in vitro* data

A high goodness of fit of the CARRGO model to the xCELLigence data was observed across all cell lines (*R*^2^ = 0.93 ± 0.1, **Fig. 3, Fig. S3**). To investigate the sensitivity of our model fitting to sampling frequency, we down-sampled the data by taking time intervals of 2 hours, 5 and 10 hours. No significant variation was observed in the model parameters *κ*_1_, *κ*_2_ and *θ* to the down-sampled data (repeated measure ANOVA p>0.1)(**Fig. S5, S6**). We consistently observed very small values of the CAR T-cell death rate (*θ* < 10^−3^). Uniqueness of the parameters was tested by choosing 100 different combinations of values of the parameters across several orders of magnitude for the model fitting optimization procedure. We found that if the optimization converged, it converged to unique values of the parameters, which is a direct consequence of the identifiability analysis of the model and minimum number of points required to resolve the model (see supplementary material1 **Fig. S7, Movie S1**).

**Figure 3:**
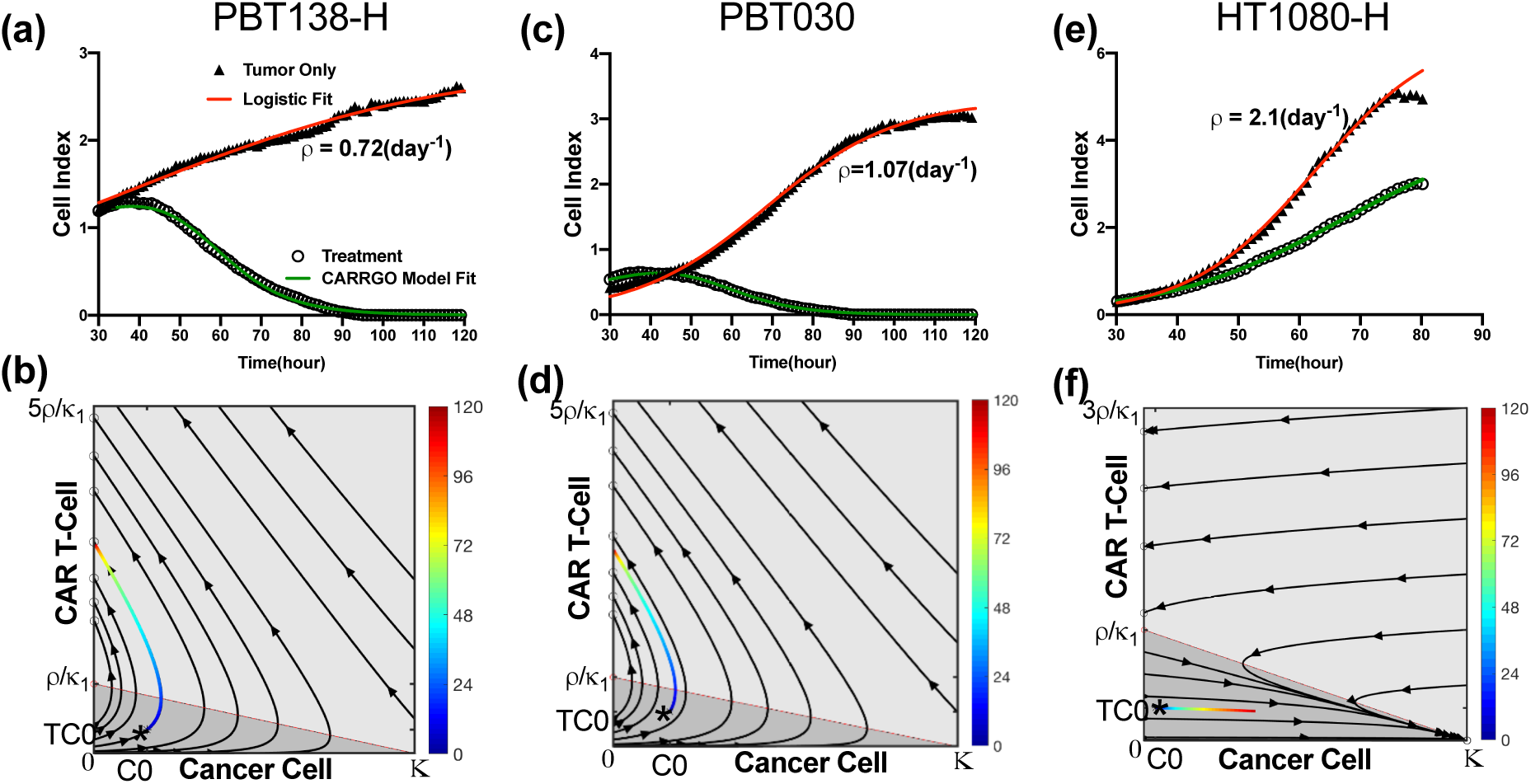
CARRGO model dynamics and in vitro CAR T-cell and glioma cell data. Dynamics from 3 cancer cell lines with and without CAR T-cell treatment along with CARRGO model fits (red line: logistic growth; green line: CARRGO model) with effector to target ratio 1:20. Top row: cell index from xCELLigence and model fit. Bottom row: data of treatment dynamics plotted in phase-space colored by time (hours). (a, b) PBT138, tumor seeding 12.5 × 10^3^ (cells), (c, d) PBT030, tumor seeding 12.5 × 10^3^ (cells), (e, f) HT1080 High, tumor seeding 2 × 10^3^ (cells). Estimated tumor growth rate for cell line PBT138-H was *ρ* = 0.72 day^−1^, for PBT030 was *ρ* = 1.07 day^−1^ and for HT1080-H was *ρ* = 2.1 day^−1^. PBT138-H and PBT030 show successful CAR T-cell killing. HT1080-H shows CAR T-cell exhaustion and treatment failure. These dynamics are captured in the phase space diagrams and CARRGO model parameters, which correctly capture transient progression before response for PBT138 and PBT030 (case 1, Figure 1) and rapid progression for HT1080 H (case 2, Figure 1).

### Validation of xCELLigence dynamics with flow cytometry

Because the xCELLigence system is an indirect measure of cell number, we validated the previously reported linear relationship [39] between the cell index read-out from the machine and the number of cells measured with flow cytometry (r^2^ > 0.9, supplementary material2 **Fig. S8**). Because cell index measures the change in electrical impedance caused by both cancer cells and CAR T-cells adhering to the plate, CAR T-cell dynamics are not directly measured by the system, so we compared the cancer cell (CC) to T-cell ratio (TC) from flow cytometry to that predicted from CARRGO model. The model predicted ratio CC/TC at end time point shows a similar trend to that measured with flow cytometry, indicating the CARRGO model-predicted CAR T-cell dynamics derived from the xCELLigence data are consistent with flow cytometry measurements. This trend was observed in PBT030 and PBT138 for BBζ and 28ζ CAR T-cells and for all doses (supplementary material2 **Fig. S9**).

### CAR T-cell dose-dependent dynamics

We examined the effect of varying the effector to target ratio, i.e. CAR T-cell dose for all cell lines. The CAR T-cell death rate parameter was found to be very small (*θ* < 10^−4^ day^−1^) for all cancer cell lines and all CAR T-cells and doses. The killing rate parameter *κ*_1_ shows a negative correlation while *κ*_2_ shows positive correlation with respect to the CAR T-cell dose. The parameter *κ*_2_ was negative only for tumor line HT1080-H which indicates the exhaustion rate being much stronger than the proliferation rate of the CAR T-cells. A positive correlation of *κ*_1_ with CAR T-cell dose indicates that higher dose of CAR T-cell results in a lower killing rate, since each individual CAR T-cell has fewer number of cancer cells to encounter. The range of values for κ_1_ and κ_2_ varied with cancer cell line as the growth rate of each cell line is different from each other, however, the overall trends were preserved across the cell lines. Plots of *θ, κ*_1_ and *κ*_2_ for PBT030, PBT138 (seeding 12.5 × 10^3^) and HT1080-H treated with BBζ CAR T-cells are shown in **Figure 4**. Other cell-lines and parameters for 28ζ CAR T-cells are given in supplementary material2 (**Fig. S10**).

**Figure 4:**
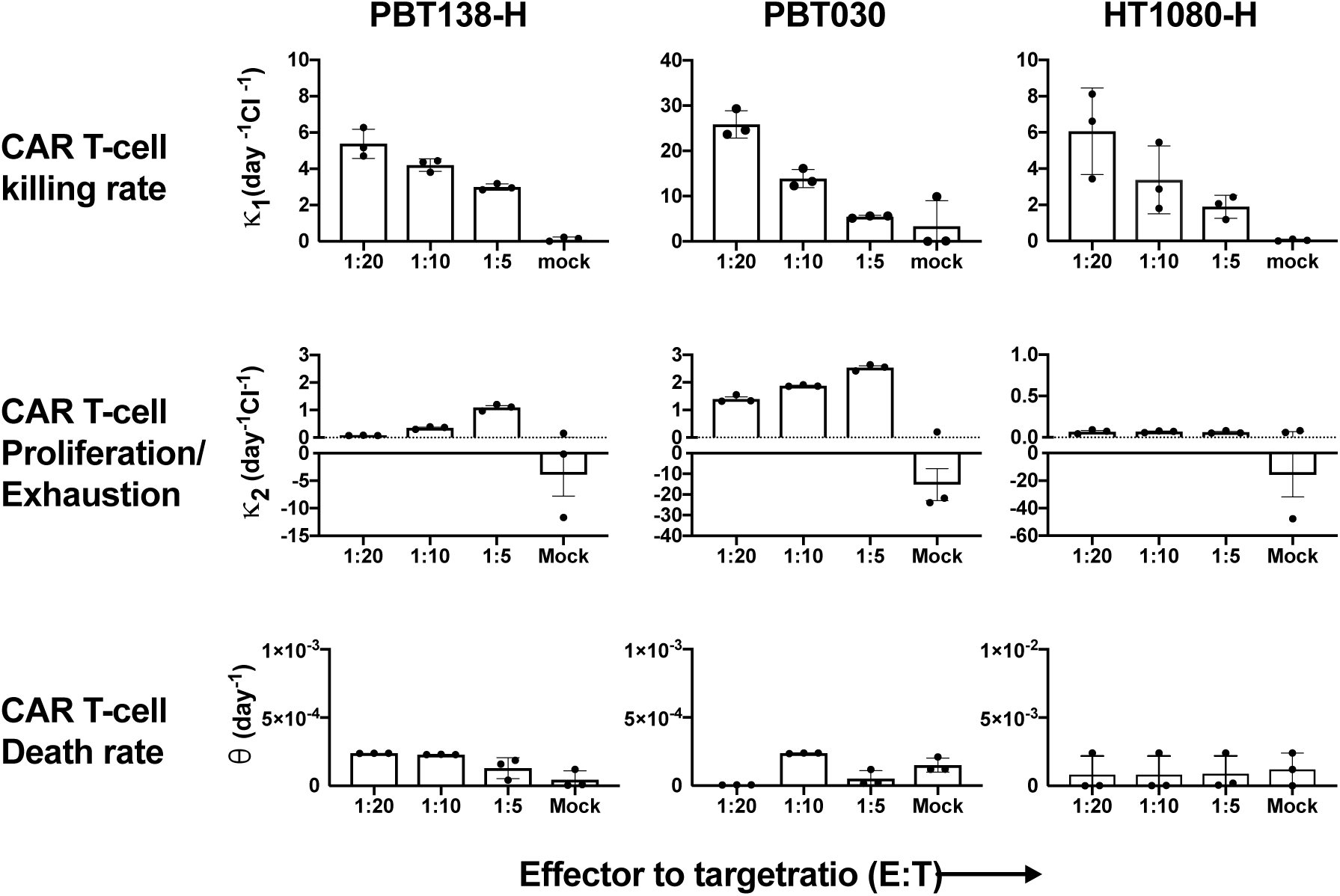
Comparisons of CARRGO model parameters with cell line and CAR T-cell dose. CARRGO model parameters: (top row) killing rate (κ_1_), (middle row) proliferation/exhaustion rate (κ_2_), (bottom row) and persistence/death rate (θ), for cell lines PBT138-H, PBT030, and HT1080-H treated with three different effectors to target (ET) ratios (1:5, 1:10, 1:20). CAR T-cell killing rate is observed to decrease with increasing ET ratio for all cell lines. This suggests CAR T-cells kill more cancer cells per unit time at a lower concentration as compared to higher ET ratio. In contrast, the CAR T-cell proliferation / exhaustion rate increases with ET ratio. This suggests that the CAR T-cells are stimulated to proliferate and are less exhausted with higher ET ratio as compared to lower. For reference, CAR T-cells are hypoactivated in mock (*κ*_2_ < 0). The CAR T-cell death rate, or persistence, is observed to be independent of target cell line and ET ratio.

### Relating *κ*_1_, *κ*_2_ with tumor growth rate and antigen expression

Tumor growth rate *ρ* varies significantly (p<0.01) among different cell-lines and with antigen expression level (see supplement material2 **Fig. S11a**). To investigate the relationship between tumor growth rate and CAR T-cell killing *κ*_1_ and exhaustion *κ*_2_, we evaluated cell lines with antigen levels greater than 80% and treated with BBζ CAR T-cells at an effector to target ratio of 1:5. No significant correlation was found between the cancer cell proliferation rate *ρ* and killing rate (*κ*_1_) (**Fig. S11b**). However, the exhaustion rate *κ*_2_ is significantly correlated with tumor growth rate (**Fig. S11c**) with Pearson correlation coefficient *r* = –0.9, p<0.001. Similar results were observed for the cells treated with 28ζ IL13Rα2-CARs. **Figure 5** shows the density of IL13R*α*2 level on cancer cell surface and its relation to CAR T-cell killing for cell line HT1080-H and PBT138-H. We observed that *κ*_1_ shows a decreasing trend from medium to high antigen level (**Fig.5c**) suggesting that high levels of antigen expression may not result in faster rates of CAR T-cell killing. The rate constant *κ*_2_ increases from low to medium antigen expression and plateaus with high levels (**Fig.5d**). This suggests limited activation of CAR T-cell at lower antigen expression and exhaustion rate from medium to high antigen may not change significantly and may be the result of over-activation of the CAR T-cells.

**Figure 5:**
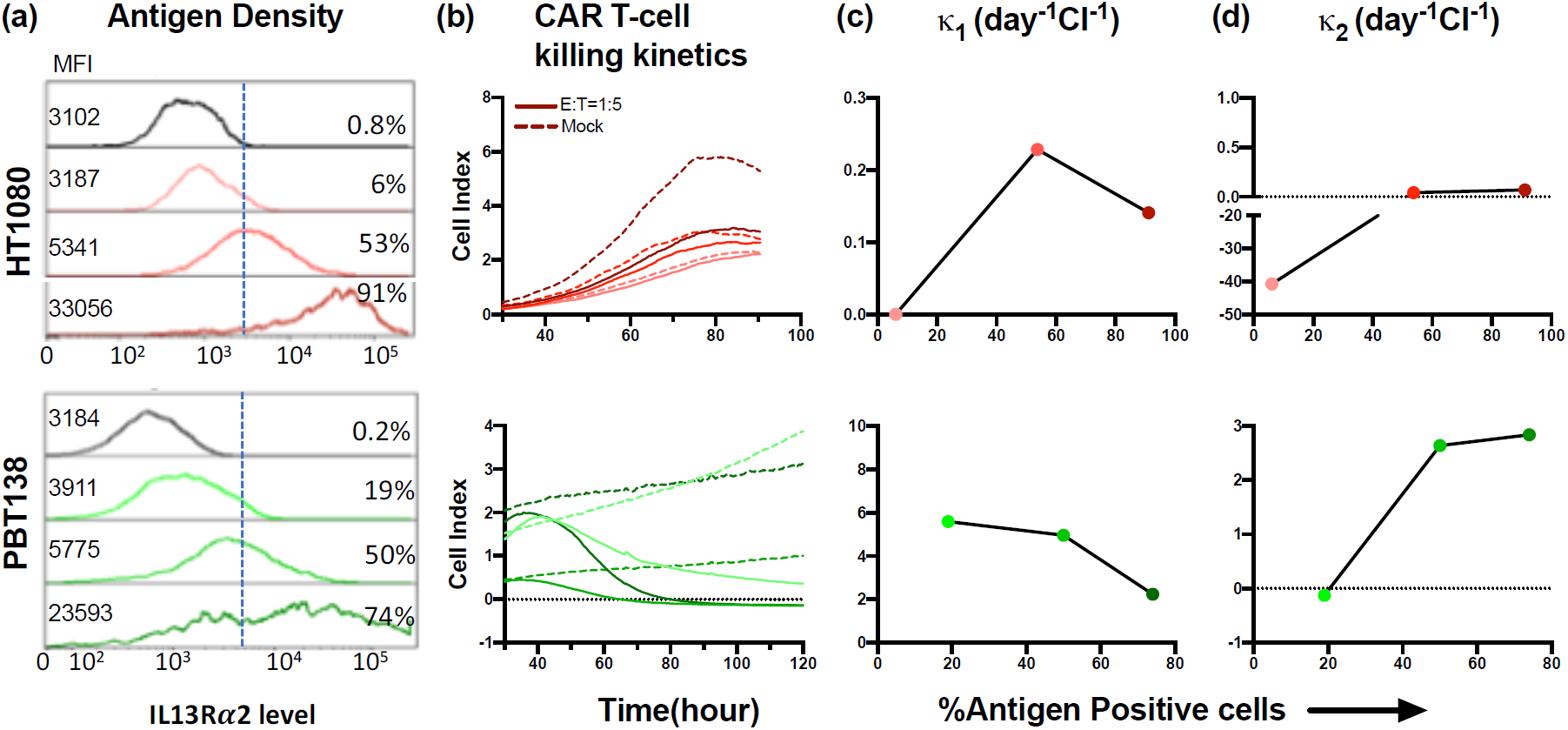
Killing kinetics of CAR T-cells as compared to antigen expression level. Antigen expression measured with flow cytometry (mean florescence intensity (MFI), % of cells positive) for cell line HT1080, PBT138 (mock, low, medium, high) (a) and CAR T-cell killing dynamics measured by xCELLigence (b). *κ*_1_ shows a decreasing trend from medium to high antigen levels (c) suggesting that high levels of antigen expression may not result in faster rates of CAR T-cell killing. *κ*_2_ increases from low to medium antigen expression and plateaus with high antigen levels(d). This suggests limited activation of CAR T-cell at lower antigen expression and that exhaustion rates from medium to high antigen may not change significantly.

### CARRGO model applied to *in vivo* human data

To translate the in vitro dynamics to of the model to real patient data [23], we fit the CARRGO model to MRI-derived tumor volume data during CAR T-cell treatment (**Fig. 6**). The CARRGO model is able to fit the tumor growth dynamics quite accurately for lesions T6, and T7 with the same set of parameters *κ*_1_ = 6 × 10^−9^ (day^−1^ cell^−1^), *κ*_2_ = 0.3 × 10^−10^ (day^−1^ cell^−1^), *θ* = 0.1 × 10^−5^ (day^−1^) and lesion T9 with *κ*_1_ = 9 × 10^−8^ (day^−1^ cell^−1^), *κ*_2_ = –2 × 10^−13^ (day^−1^ cell^−1^), *θ* = 5 × 10^−5^ (day^−1^). In the case of lesion T9, although the CARRGO model is consistent with the overall tumor dynamics, it does not fit the later time points following CAR T-cell treatment well. This is because lesion T9 received radiation treatment between day 200 to 300, which is not included in the CARRGO model. We note the negative correlation between the tumor growth rate (*ρ* = 0.06, 0.07 /day and *ρ* = 0.2 /day for T6, T7 and T9 respectively) with the CAR T-cell exhaustion rate *κ*_2_ in the patient data, which is consistent with that observed in the experimental data (**Fig. S11c**). We remark that the parameters *κ*_1_ and *κ*_2_ are on the order of *O*(10^−13^), which appear to be very small, however, these parameters are scaled by the caring capacity *K* in units of cells, which is of order *O*(10^9^). Therefore, these parameter values are comparable with the in vitro data when scaled relative to the carrying capacity (**Fig. 4**)

**Figure 6:**
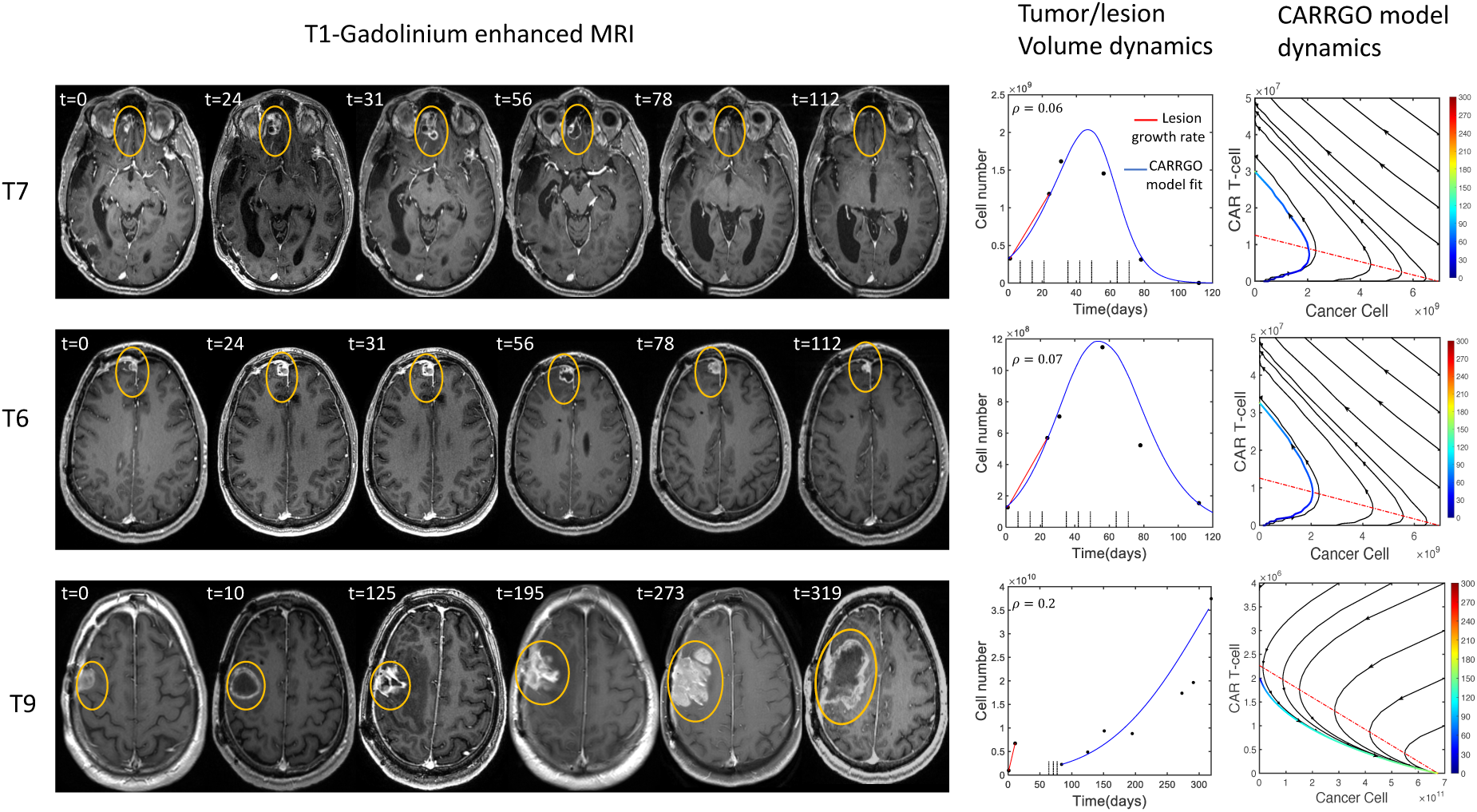
CARRGO model applied to in vivo human data. A male patient with multi-focal glioblastoma was treated with IL13Rα2 CAR T-cells. Yellow circles are used to indicate tumor location and do not reflect tumor size. Cell number is calculated from tumor size with volumetric segmentation of the contrastenahcing lesion. The right columns shows CARRGO model fits and dynamics based on the tumor volume data. The CARRGO model parameters are the same for lesions T7 and T6, which responded to CAR T-cell treatment. The model predicts that the non-responding lesion T9 had a smaller rate of CAR T-cell killing and increased rates of exhaustion and CAR T-cell death. Lesion T9 was also observed to have a higher cancer cell proliferation rate (*ρ* =0.2/day) as compared to T6 and T7 which had very similar rates (*ρ* =0.06/day, *ρ* =0.07/day, respectively).

## Discussion

The CARRGO model is a simple representation of cancer-cell T-cell interactions. We developed the CARRGO model with the aim of understanding CAR T-cell efficacy in terms of rates of killing, proliferation, exhaustion, and persistence with a real-time cell analyzer in a simple, controlled *in vitro* system. The CARRGO model fit remarkably well to the highly temporally resolved experimental data and as well as to data derived from a patient treated with IL13Rα2 BBζ CAR T-cells. Although the predator-prey mathematical model formalism has been widely used in a number of biological settings, the novelty of this model is in the application to a novel form of cancer therapy with a high temporal resolution cell monitoring experimental design which provides nearly continuous data on killing kinetics.

With the CARRGO model we show that the rate of cancer cell killing by CAR T-cells is inversely related to the CAR T-cell dose. With a fixed number of cancer cells as an initial condition, as the number of CAR T-cells increases (dose), any individual T-cell encounter will encounter fewer number of cancer cells to kill, indicating that increasing dose does not result in a maximal rate of killing on a per T-cell basis. For example, the PBT138 cell line shows complete killing within 80 and 100 hours for effector to target ratios 1:5 and 1:10 respectively. This result suggests that a lower dose of CAR T-cells may change the time to complete cancer cell killing but shows the same overall cancer cell killing effectiveness. We observed that *κ*_2_ positively correlated with CAR T-cell dose. Because the parameter *κ*_2_ is a net measure of CAR T-cell proliferation and exhaustion or lack of activation and the T-cell proliferation is not dose dependent [18], the trend observed in *κ*_2_ with dose is dominated by the exhaustion rate: the higher the dose, the lower the exhaustion rate, resulting in an increased value of *κ*_2_. The death rate of CAR T-cells was very small as compared to the cancer cell proliferation rate for all conditions. This is likely due to the short time-scale of the experiment and because the T-cells were stimulated to proliferate by the presence of cancer cells.

We observed that the cancer cell growth showed no relation with CAR T-cell killing rate and an inverse relationship with *κ*_2_. This may explain variations in patient-specific responses even for the same CAR T-cell dose. For a fixed CAR T-dose, *κ*_2_ is the principle determinant of treatment failure or success as shown in phase plane analysis (**Fig.1**), which is also observed in patient data (**Fig.6**). This result, driven by the CARRGO model analysis suggests that the balance between proliferation and exhaustion of CAR T-cells may contribute more than the rate of CAR T-cell mediated cancer cell killing in determining treatment success or failure. Moreover, the CARRGO model predicts transient progression of cancer cells even in the case of successful CAR T-cell therapy. This prediction may be consistent with the clinical phenomenon of pseudo-progression, in which the cancer is seen to progress during therapy before eventually responding [7,8]. Identifying characteristics of the patient and the CAR T-cells which may result in pseudo-progression could have a profound effect on interpretation of these dynamics observed in the clinic.

Interestingly, we found *κ*_1_ decreases and *κ*_2_ plateaued from medium antigen level to higher level of antigen expression. One of the possible explanations of this behavior could be the antigen density is more heterogeneous in the higher antigen level cell population as compared to medium and low antigen levels (**Fig.5a**). A ore heterogeneity in the density of antigen expression intensity in the cancer cells within the initial population may cause clustering of CAR T-cells resulting in their exhaustion [20,41]. Another confounding factor can be the dependence of the detected antigen signal intensity on both the number of antigen-positive tumor cells and their individual antigen expression intensity. Elucidating these factors individually can better tune the model parameters and the prediction of the tumor response dynamics. However, more studies are required to examine effect of cell and population level antigen density on CAR T-cell killing kinetics.

There are some important limitations to consider with this model and experimental system. Perhaps the most obvious is that the *in vitro* system is not a model for the human immune system or tumor microenvironment. It does not include cytokines, stromal cells, or additional immune cells such as myeloid cells which contribute to CAR T-cell activity in vivo. Another limitation is the assumption that the populations are well mixed. In practice, this assumption may depend on the route of CAR T-cell administration, with intracavitary and intraventricular injections potentially resulting in spatially heterogeneous densities of CAR T or cancer cells, although methods to assess the distribution of CAR T-cells in vivo remains an open challenge[42,43]. To address this limitation, the well-mixed assumption may be relaxed and CAR T-cell killing dynamics interrogated with spatial or agent-based models[44]. Another limitation is with regard to the experimental system: the change in electrical impedance measured by the cell index does not differentiate cell detachment from cell killing. This is only a minor consideration as the cell lines used are very adherent to the plate and were not observed to detach. Finally, the experimental system does not directly measure dynamics of CAR T-cells. However, our model is initialized with known numbers of cancer cells and CAR T-cells and the model-predicted cancer cell to CAR T-cell ratio at the experimental endpoint was validated with flow cytometry, giving confidence to our model predictions and parameter estimates. To address this limitation, the CARRGO model CAR T-cell dynamics can be validated by labeling the CAR T-cells and directly measuring their dynamics with live cell imaging based methods [45]. Despite these limitations, the CARRGO model succeeded in revealing nonlinear dynamics, quantifying kinetics of killing, and generating hypotheses which may be tested in other *in vitro* systems, and other computational or *in vivo* models.

In summary, CAR T-cells have shown promise in hematologic malignancies and are being actively investigated in solid tumors. We aimed to use mathematical modeling to investigate factors which contribute to the kinetics of CAR T-cell mediated cancer cell killing in a simple isolated *in vitro* system. We were able to fit the CARRGO model to *in vitro* and *in vivo* human data with remarkable accuracy. We demonstrated that we can consistently and reproducibly estimate rate constants in the CARRGO model and investigate their dependence on CAR T-cell dose and antigen expression levels. The CARRGO model may be combined with other mathematical models which estimate cancer cell growth and proliferation rates noninvasively with MRI data [9,11,46] to produce a fine-tuned and benchmarked suite of mathematical models, which may aide in optimization of dosing and scheduling of CAR T-cells for greater individualized and personalized therapy.

## Acknowledgements

Research reported in this publication was supported by the California Institute for Regenerative Medicine (CLIN2-10248), the National Cancer Institute of the National Institutes of Health under grant numbers R01CA236500, P30CA033572. The content is solely the responsibility of the authors and does not necessarily represent the official views of the California Institute for Regenerative Medicine or National Institutes of Health.

