## Supplementary material for "Mathematical deconvolution of CAR T-cell proliferation and exhaustion from real-time killing assay data": Table S1 (from another version of this preprint: version 1, 2019-09-30)

**Cells and cell culture**

Three cell lines were used in this study PBT030, PBT138, HT1080. All cell lines were positive for IL13R$\alpha$2 and antigen level >80%. PBT138 and HT1080 was lentivirally modified to express varied levels of IL13Rα2. Initially PBT030 and PBT138 were seeded with both 12.5k and 25k cells to investigate effect of cell seeding on tumor cell proliferation. Later 12.5K were decided to seed to keep the system below confluency for longer time period. As HT1080 is a highly proliferative cell line, only 2K number of cells were seeded to keep the system below confluency.

**Table S1:** Cancer cell lines, CAR T-cells, seeding and effector to target ratios used in the *in vitro* experiments.

| Tumor Cell Line | CAR T-cell | Tumor cell seeding | Effector : Target |
|---|---|---|---|
| PBT030 | BB$\zeta$, 28$\zeta$ | 12.5K, 25K | 1:20, 1:10,1:5 |
| PBT138-High | BB$\zeta$, 28$\zeta$ | 12.5K, 25K, | 1:20, 1:10, 1:5 |
| HT1080-High, HT1080-Medium, HT1080-Low | BB$\zeta$, 28$\zeta$ | 2K | 1:20, 1:10, 1:5 |
| PBT138-High, PBT138-Medium, PBT138-Low | BB$\zeta$, 28$\zeta$ | 12.5K | 1:20, 1:10, 1:5 |
