## Supplementary material for "Mathematical deconvolution of CAR T-cell proliferation and exhaustion from real-time killing assay data": Fig S1-S7 (from another version of this preprint: version 1, 2019-09-30)

**SUPPLEMENTARY DATA 1 (Figs. S1-S7)**
**Data supporting CARRGO model assumptions**

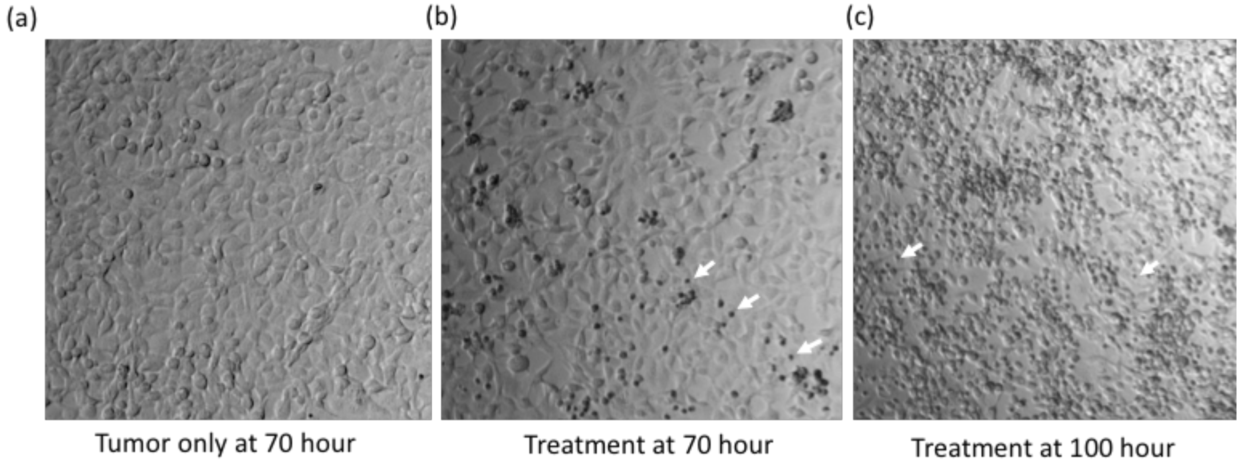


**Figure S1. Well-mixed population.** Microscopic images of HT1080-High IL13Rα2-expressing tumor only (a) at 70 hours post seeding during Logistic growth, and cancer cells with CAR T-cells (b) at 70 hours and (c) at 100 hours post seeding. White arrows indicate examples of CAR T-cells which appear as small dots as compared to the larger cancer cells. CAR T-cells are well-mixed with the cancer cell population. Both cancer cells and CAR T-cells are spread/attached on the bottom of the well. Because the CAR T-cells are small as compared to cancer cells, they contribute very little to the measured cell index.

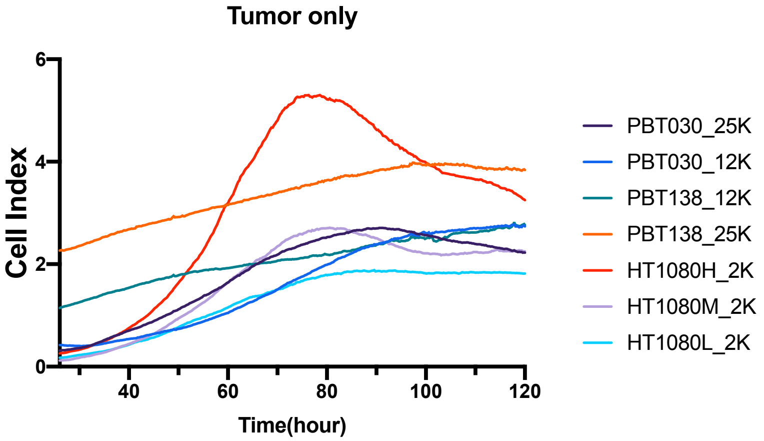


**Figure S2. Logistic growth**. Cancer cell dynamics of different cell lines at various seeding densities demonstrate logistic growth. Cancer cell lines shown here are only from the untreated wells and demonstrate that long-term growth dynamics are limited by physical and nutrient constraints of the system. We note that HT1080H,M reach confluency at approximately 80 hours post seeding. The linear relation between CI and cell number is lost after the cells reach confluency therefore we used only up to 80% of the confluency data in our CARRGO model fitting (c.f. Figure 2). PBT lines reached confluency at approximately 120 hours post seeding.

### Raw xCELLigence data and CARRGO model fits

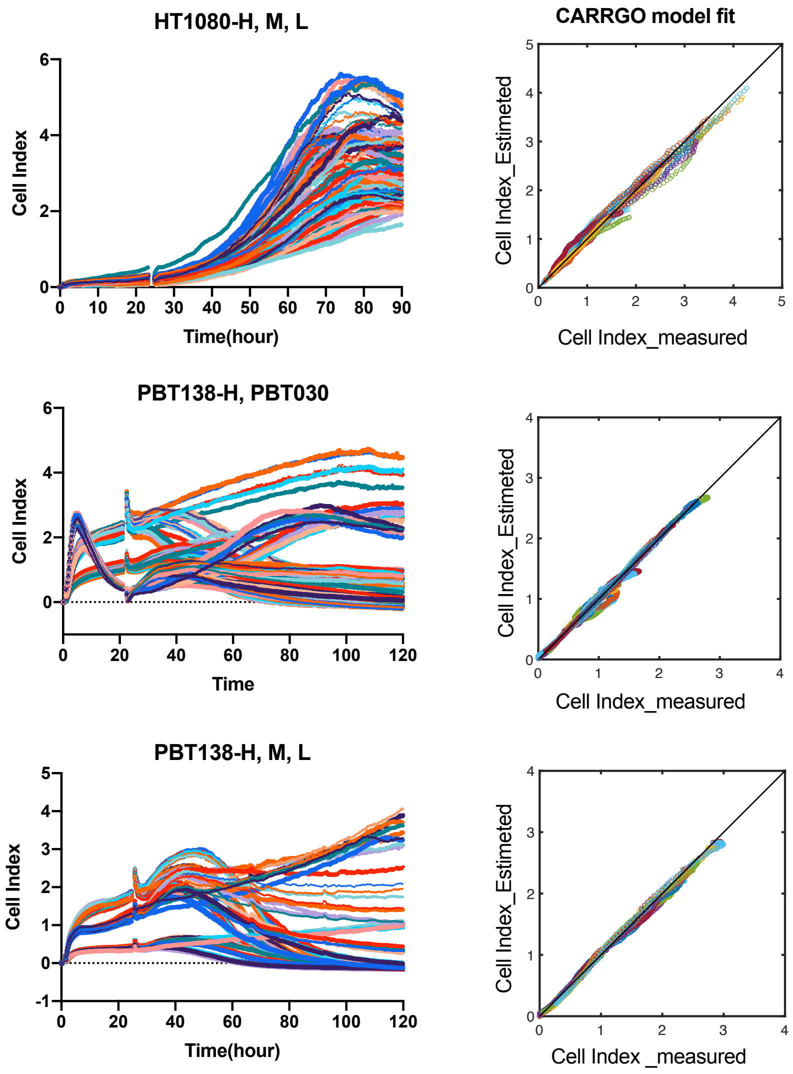


**Figure S3**. **CAR T cell killing kinetics as measured by xCELLigence.** (Left column) IL13Rα2 CAR T-cells and cancer cell lines with varying IL13Rα2 antigen expression showing cancer cell dynamics in the 96 well E-plate measured by xCELLigence. (a) HT1080-IL13Rα2 lines engineered for low, med, and high receptor expression. (b) PBT030, a glioma line that endogenously expresses IL13Rα2, and PBT138 a glioma line that does not express IL13Rα2 endogenously (c) PBT138-IL13Rα2 lines engineered for low, med and high receptor expression. Cells were either left untreated (triplicates per cell line) or treated with CAR T-cells with effector to target ratios of 1:5, 1:10, and 1:20. Each cell line was treated with three CAR T-cells: BBζ, 28ζ, and mock. (Right column) Correlation between measured data (X-axis) and CARRGO model estimated data (Y-axis) with $R^2$=0.9±0.1 (middle row; PBT138-H, PBT030). Data before 24 hours is attachment and spreading phase and not included in the CARRGO model fitting (c.f. Figure 2). Each color represents data from a single well in a 96 well plate (rows).

### Partial differential equation model of cell attachment and spreading

We assume that the attachment of cell into the bottom of the well is a diffusion process. **Reaction–diffusion systems** are mathematical models which correspond to change of concentration of substances in space and time. Change in concentration of cancer cell population (C) can be written as

Where $D$ is the diffusion coefficient (mm$^2$/time), $\rho$ is the net growth rate (1/time) of cancer cells, $K$ is the cancer cell carrying capacity (cells or CI).

$$\overbrace{\frac{\partial C}{\partial t}}^{\substack{rate\ of\ change\ of \\ cancer\ cell\ population}} = \overbrace{\frac{\partial}{\partial x}\cdot\left(D\frac{\partial}{\partial x}C\right)}^{\substack{Dispersion\ of\ cells\ on \\ bottom\ of\ the\ well}} + \overbrace{\rho C\left(1-\frac{C}{K}\right)}^{Logistic\ growth}$$

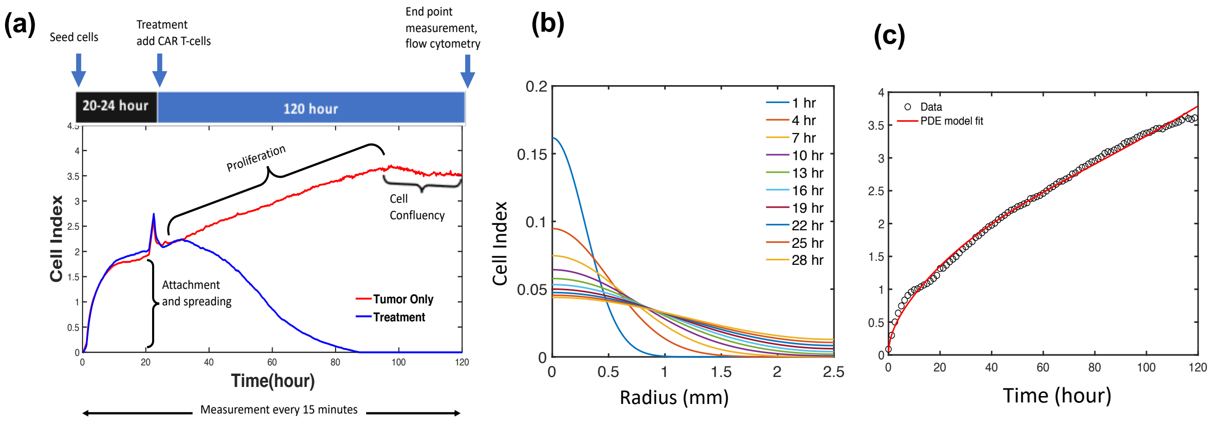


**Figure S4. Partial differential equation model of attachment and spreading in xCELLigence system.** Initial phase of cancer cell dynamics in the experimental system (a), explaining the attachment and spreading of the cells on the bottom of the plate. (b) solutions of the PDE model showing spatial distribution of cells over time with a point-source (Dirac delta function centered at x=0) initial condition at time t=0, (c) Reaction-diffusion model fitting to the tumor only growth curve data. A diffusion rate D = 0.01 mm$^2$/hour was used in fitting the reaction-diffusion PDE model to the data. The PDE model fits the attachment and spreading dynamics remarkably well. Because the cell populations are well mixed (Figure S1) we neglect the spatial term in the CARRGO model once the diffusion process is near steady state ($\frac{\partial^2 C}{\partial x^2} \approx 0$, t ~ 24 hours). We show the PDE model fit to this early attachment and spreading phase data for the reader's interest and information.

**Sensitivity of CARRGO model fitting to temporal frequency**

To investigate the stability of the CARRGO model parameters, we varied the temporal resolution of the data by down-sampling. Data was down-sampled by taking data points in time intervals ranging from 15 minutes (full data) to 10 hours. Top row shows the value of the parameters $\kappa_1$, $\kappa_2$ and $\theta$ for all down-sampled data of a single well. $\kappa_1 = 0.18 \pm 0.003$, $\kappa_2 = 0.08 \pm 0.0005$ and $\theta = 0.001 \pm 2.5 \times 10^{-6}$. Repeated measure ANOVA was performed to check the variability in the parameter values at 1 hour, 5 hours, and 10 hours of down-sampled data. No significant difference (p>0.1) was found in the ANOVA test. Bottom row shows the fitting of the model to the data for 1, 5, and 10-hour time intervals. The CARRGO model fit well to all down-sampled data with $R^2 \approx 0.9$, indicating consistency and reproducibility of the model and the data.

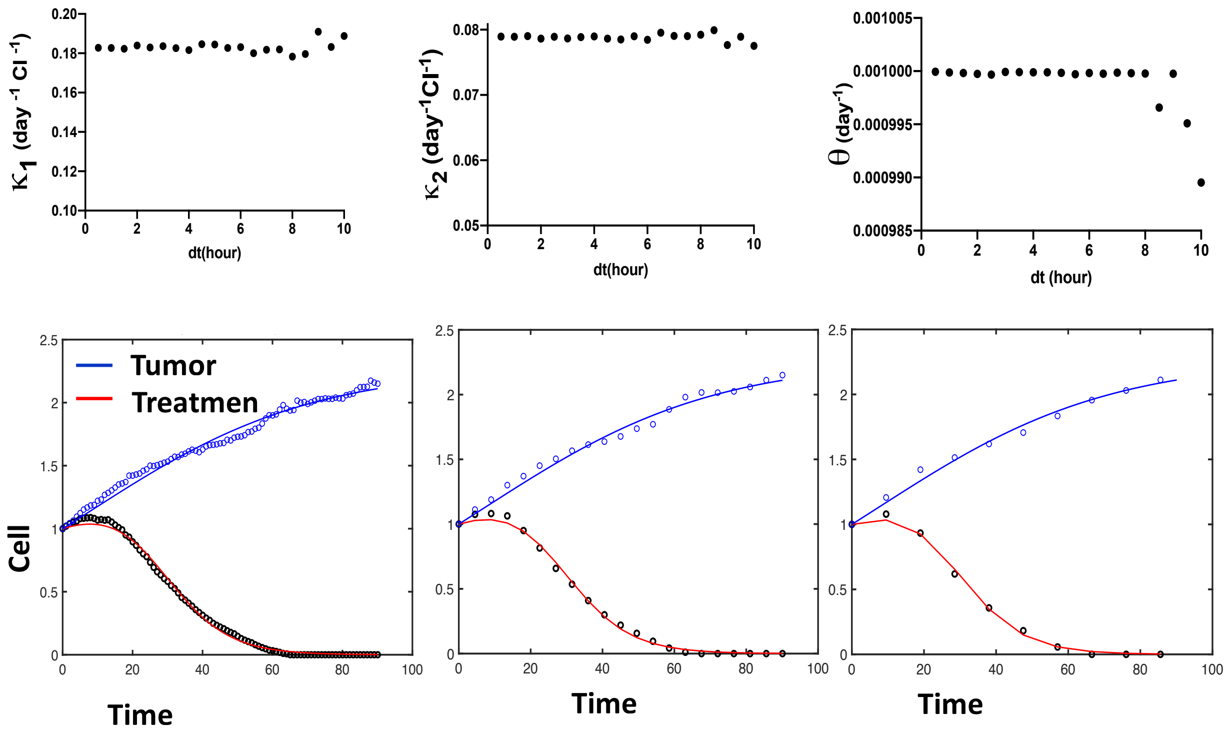


**Figure S5. Effect of data sampling on CARRGO model parameter estimates.** Top row shows the value of the parameters $\kappa_1$, $\kappa_2$ and $\theta$ for all down-sampled data of a single well for cell line PBT138-H treated at dose E:T=1:5. Bottom row shows the fitting of the model to the data of 1, 5, and 10 hour time intervals.

### Sensitivity of CARRGO model fitting to the end time of measured data point

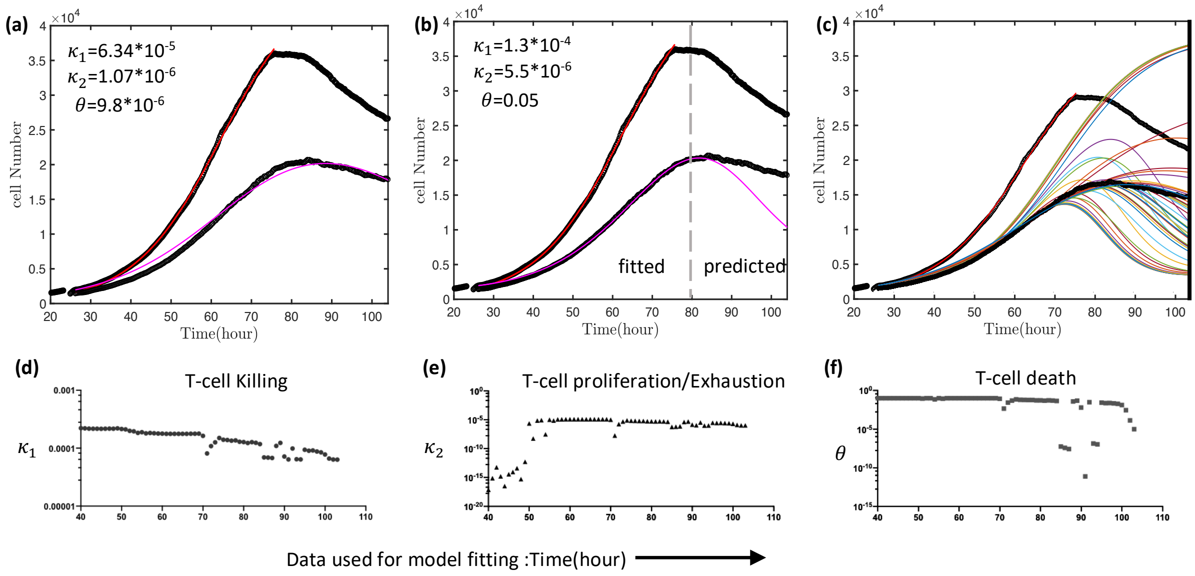


**Figure S6. Sensitivity of CARRGO model fitting and prediction to the end time of measured data point.** (a) CARRGO model fitting when the full time series of data was used. (b) CARRGO model fitted for only for 80 hour and dynamics for later time points are predicted. (c) The model predicted dynamics when fewer time series data are used for model fitting and corresponding $\kappa_1$, $\kappa_2$, $\theta$ values are shown in (d), (e), (f) respectively. This suggests $\kappa_2$ is the most sensitive parameter to predict the tumor dynamics.

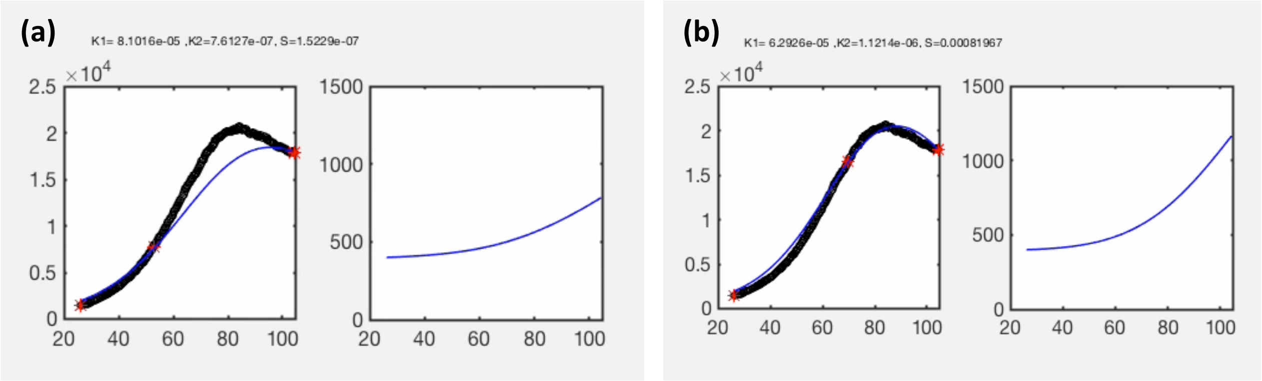


**Movie S1. CARRGO model fit to only three data points.** Because there are three treatment-related parameters in the CARRGO model, three is the minimum number of data points needed to identify these parameters. We examined the sensitivity of parameter identification by using the initial (t=24 hours post seeding) and final time points and varied the third time point. (Left plot) Black points are raw xCELLigence data for HT1080-H treated with 1:5 effector to target ratio BBζ CAR T-cells. Red asterisks are data used to fit CARRGO model. Blue curve is CARRGO model solution. (Right plot) CARRGO predicted CAR T-cell population. (a) Third data point at t = 50 hours. (b) Third data point at t = 70 hours. The best fit to the whole data occurs when the third point lies between the growth curve inflection point (t~65 hours) and maximum (t~80 hours). We include a movie which shows the effect of varying the third data point from t = 24 to t = 110 hours.

### Simulation to test uniqueness of CARRGO model parameters

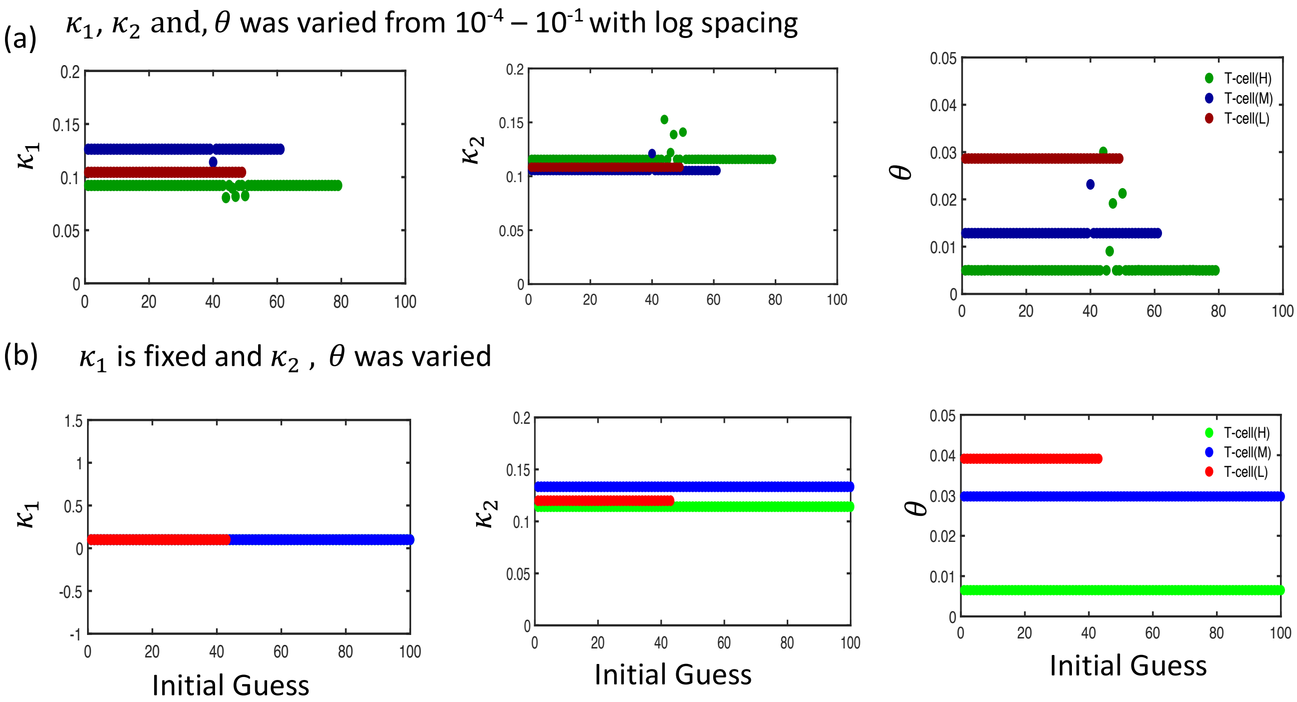


**Figure S7. Simulation to test uniqueness of CARRGO model parameters.** Uniqueness of the parameters was tested by choosing 100 different combinations of values of the parameters across several orders of magnitude for the model fitting optimization procedure. We found that if the optimization converged, it converged to unique values of the parameters. Top row shows the converged parameter values started from 100 different combination initial guess in parameter ranges from $10^{-4}$ – $10^{-1}$ with log spacing and 300 iterations. Except for a few points, the model converges to unique set of parameters. Those points again converge to the unique solution when number of iterations is increased. Bottom row shows the convergence of the model when one of the parameters is fixed. With a constrained variable, the optimization converged to a unique set of parameters with fewer iterations. Since the value for $\theta$ was very small, we constrained $\theta$ for all CARRGO data fitting.
