## Supplementary material for "Mathematical deconvolution of CAR T-cell proliferation and exhaustion from real-time killing assay data": Fig S8-S11 (from another version of this preprint: version 1, 2019-09-30)

**SUPPLEMENTARY DATA 2 (Figs. S8-S11)**

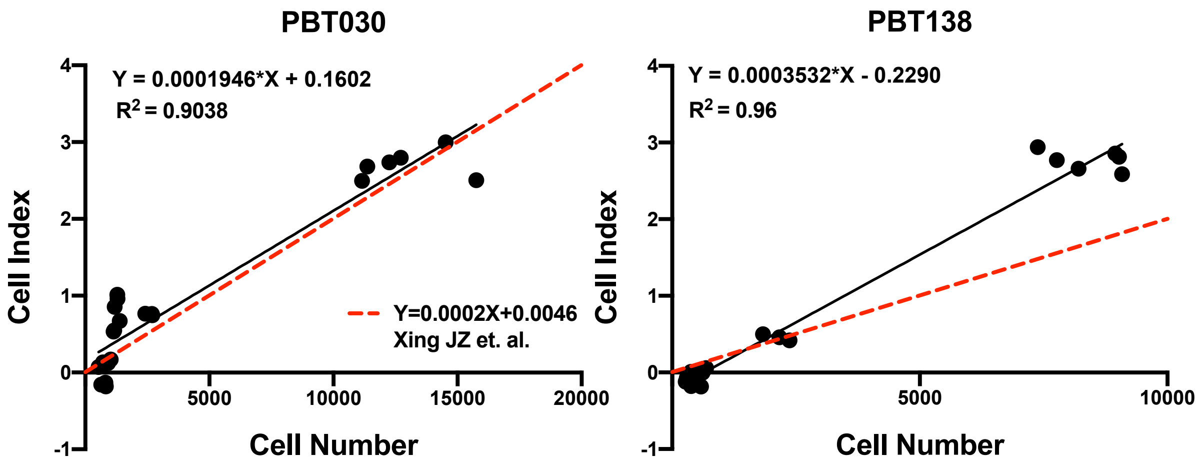


**Figure S8. Linear relation between cell index (CI) and cell number (CN).** A strong linear relationship exists between the xCELLigence output of cell Index and the total number of cells in the well. Data is shown for cell lines PBT030 and PBT138-H. Cell numbers were measured with flow cytometry at 120 hours after cell seeding at the end of the experiment. Measurements were taken from all wells in the 96 well plate with seeding densities 12.5k. The red line shows the slope of published report by Xing et al. (1).

Because the xCELLigence system is an indirect measure of cell number, we validated previously published relationships between cell number and the output of the xCELLigence system, Cell Index (CI) by performing flow cytometry at the end of the experiments. The strong correlation ($R^2 > 0.90$) and slope is consistent with previously published reports (1). The linear relation between cell index and cell number has been confirmed by many authors, however, the slope is very much cell line dependent(1,2).

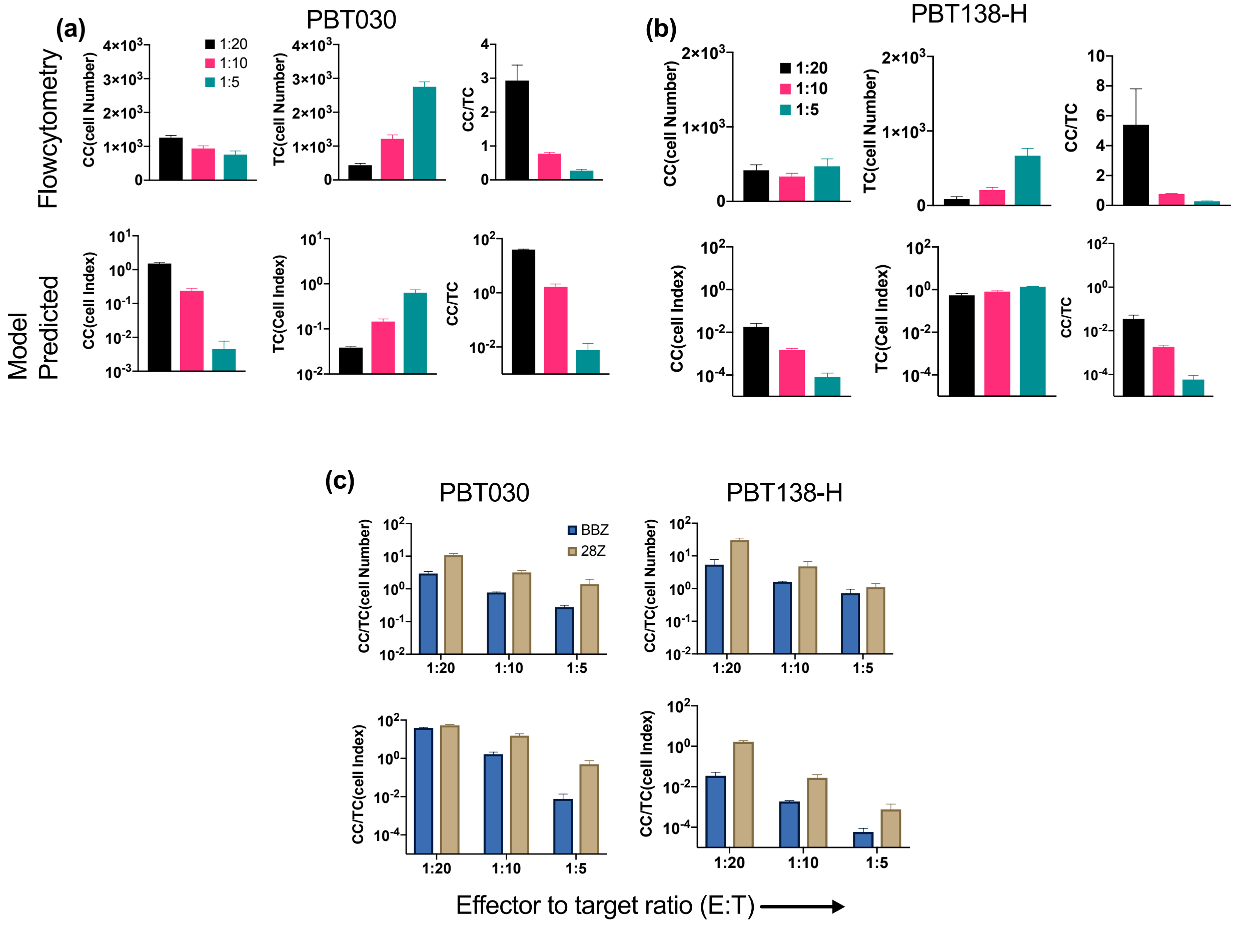


**Figure S9. Validation of CARRGO predicted cancer cell - CAR T-cell ratio by flow cytometry.** Bar plots (a), (b) showing comparison between flow cytometry (top row) and model predicted (bottom row) cancer cell (CC), CAR T-cell (TC) and the ratio CC/TC for cell lines PBT030 and PBT138-H respectively, treated with BB$\zeta$ CAR T-cells. (c) Bar plot shows cancer cell to T-cell ratio for both cell lines and comparison of BB$\zeta$ and 28$\zeta$. Note the decreasing trend of CC/TC with respect to CAR T-cell dose observed in flow cytometry is preserved in model predicted CC/TC. Similar trends were observed for both type of CAR T-cells BB$\zeta$ and 28$\zeta$. However, CC/TC for the cells treated with 28$\zeta$ is higher than BB$\zeta$ indicating BB$\zeta$ may present superior treatment outcome than 28$\zeta$. This result is consistent with previously published literature (3,4).

Because flow cytometry and the xCELLigence systems are inherently different measurements with different units, we cannot compare them directly. Rather, we observe trends across target to effector ratios and cell lines to validate the CARRGO model predictions.

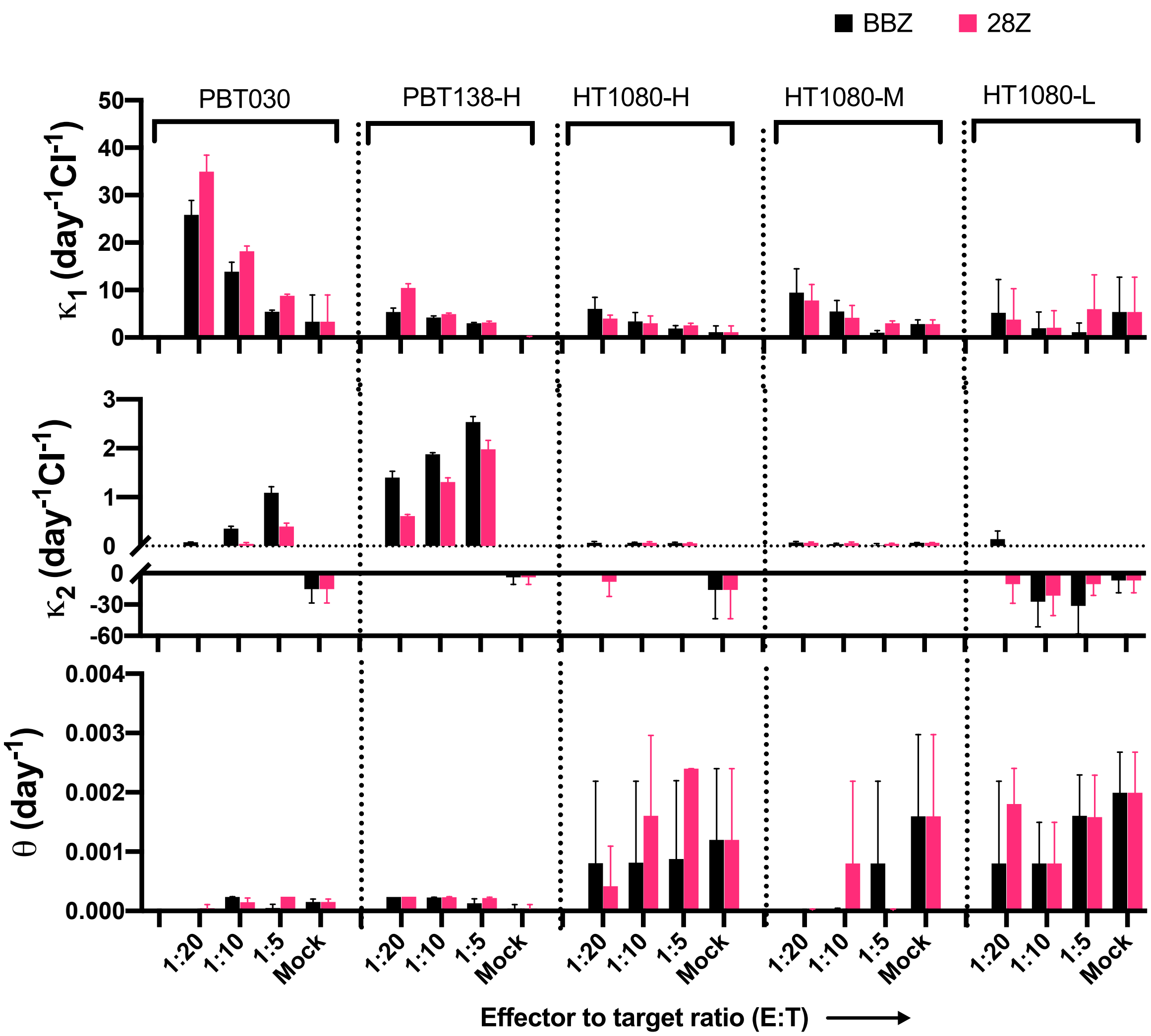


**Figure S10**. **CARRGO model parameters for different cell lines and comparison between BBζ and 28ζ IL13Rα2-CAR T-cells.** (top row) Bar plots showing killing rate ($\kappa_1$), (middle row) exhaustion rate (i.e., loss of effector activity) ($\kappa_2$), (bottom row) and net death rate ($\theta$), of PBT030, PBT138-H, and HT1080-H, HT1080-M, HT1080-L with tumor seeding $12.5 \times 10^{-3}$, three different effectors to target ratios (1:5, 1:10, 1:20). Black bars are are parameters for cells treated with CAR T-cell BBζ and pink bars for cells treated with 28ζ. $\kappa_1$ and $\kappa_2$ shows similar trend for both CAR T-cells BBζ and 28 ζ.
