## Supplement method for "Mathematical deconvolution of CAR T-cell proliferation and exhaustion from real-time killing assay data" (from another version of this preprint: version 1, 2019-09-30)

### STRUCTURAL IDENTIFIABILITY ANALYSIS OF CARRGO MODEL

$$\frac{dX}{dt} = \rho X(1 - \frac{X}{K}) - \kappa_1 XY \tag{1}$$

$$\frac{dY}{dt} = \kappa_2 XY - \theta Y \tag{2}$$

Let us examine the structural identifiability of the model with unknown parameters $\{\kappa_1, \kappa_2, \theta\}$ and observable quantities $\{X(t,p) = X, Y(t,p) = Y\}$. We assume that the parameters $\rho$ and $K$ are known as well as the initial condition values. We will attempt to compute this using Taylor series approach and generating series approach.

The Taylor series approach assumes that the observation functions are analytic in a neighborhood of $t = 0$, and the their successive time derivatives are measurable and contain information about the parameters to be identified. Using the measurable quantities at the successive time as $t = 0^+$. The problem reduces to determining the number of solutions for the parameters in a set of algebraic equations. From the system equations, we have

$$X'(0^+) = \rho X(0^+)(1 - \frac{X(0^+)}{K}) - \kappa_1 X(0^+)Y(0^+)$$
$$Y'(0^+) = \kappa_2 X(0^+)Y(0^+) - \theta Y(0^+).$$

Be denoting the measurable initial quantities as $X_0 = X(0^+)$, $X_1 = X'(0^+)$, $Y_0 = Y(0^+)$, $Y_1 = Y'(0^+)$, we have

$$X_1 = \rho X_0(1 - \frac{X_0}{K}) - \kappa_1 X_0 Y_0,$$
$$\kappa_1 = \frac{\rho}{Y_0}(1 - \frac{X_0}{K}) - \frac{X_1}{X_0 Y_0},$$

so that $\kappa_1$ is structurally identifiable. For the other parameters, we consider an additional derivative as $Y'' = \kappa_2 X'Y + \kappa_2 XY' - \theta Y'$, and measurable $Y_2 = Y''(0^+)$. We have

$$Y_1 = \kappa_2 X_0 Y_0 - \theta Y_0$$
$$Y_2 = \kappa_2(X_1 Y_0 + X_0 Y_1) - \theta Y_1,$$

so that we can solve for the remaining parameters as

$$\kappa_2 = \frac{-Y_1^2 + Y_0 Y_2}{X_1 Y_0^2}, \qquad \theta = \frac{X_0 Y_0 Y_2 - X_1 Y_0 Y_1 - X_0 Y_1^2}{X_1 Y_0^2}.$$

*Date*: September 27, 2019.

Therefore, the variables $\{\kappa_1, \kappa_2, \theta\}$ are structurally identifiable except at most a set of points of zero measure, that is, points for $X_0 = 0, Y_0 = 0, X_1 = 0$. In addition, the generating series approach using the MATLAB library GENSSI computes three Lie derivatives and verifies that $\{\kappa_1, \kappa_2, \theta\}$ are structurally locally identifiable.
